# SIBR-Cas enables host-independent and universal CRISPR genome engineering in bacteria

**DOI:** 10.1101/2021.04.26.441145

**Authors:** Constantinos Patinios, Sjoerd C.A. Creutzburg, Adini Q. Arifah, Belén A. Pérez, Colin J. Ingham, Servé W.M. Kengen, John van der Oost, Raymond H.J. Staals

## Abstract

CRISPR-Cas is a powerful tool for genome editing in bacteria. However, its efficacy is dependent on host factors (such as DNA repair pathways) and/or exogenous expression of recombinases. In this study, we mitigated these constraints by developing a simple and universal genome engineering tool for bacteria which we termed SIBR-Cas (Self-splicing Intron-Based Riboswitch-Cas). SIBR-Cas was generated from a mutant library of the theophylline-dependent self-splicing T4 *td* intron that allows for universal and inducible control over CRISPR-Cas counterselection. This control delays CRISPR-Cas counterselection, granting more time for the editing event (e.g., by homologous recombination) to occur. Without the use of exogenous recombinases, SIBR-Cas was successfully applied to knock-out several genes in three bacteria with poor homologous recombination systems. Compared to other genome engineering tools, SIBR-Cas is simple, tightly regulated and widely applicable for most (non-model) bacteria. Furthermore, we propose that SIBR can have a wider application as a universal gene expression and gene regulation control mechanism for any gene or RNA of interest in bacteria.

## INTRODUCTION

Homologous recombination (HR) combined with CRISPR-Cas counterselection is a powerful approach for genome editing in a wide range of bacterial species^1^. However, to achieve high editing efficiencies, generation of the desired edit through HR should precede CRISPR-Cas counterselection. Consequently, for organisms where the frequency of HR is low, genome editing through CRISPR-Cas counterselection may be cumbersome.

To enhance HR frequencies, the heterologous expression of recombinases has been used. However, this method is often laborious (maintenance of multiple plasmids) and success is not guaranteed as the recombinases may be incompatible with the target organism^2^. As an alternative, several regulation systems have been developed to control the expression and activity of the CRISPR-Cas module. Examples include the use of inducible promoters, inducible intein splicing^3^, split Cas proteins^4, 5^, inducible conformation change^6^, inducible inhibition through aptamers^7^ and inducible translation through riboswitches^8^. Other approaches focused on the inducible guide RNA functionality using ribozymes^9^, riboswitches^10, 11^ and photocaging^12^. The control of the CRISPR-Cas modules gives enough time for HR to occur before CRISPR-Cas counterselection is induced. Whilst the existing approaches are suitable for the organism, Cas protein or gRNA of interest, these solutions are typically not widely applicable. Therefore, CRISPR-Cas engineering tools would benefit from universal and tight regulation of their counter selective properties to provide enough time for HR to take place.

To address this issue, we developed the Self-splicing Intron-Based Riboswitch (SIBR) system. SIBR is based on the bacteriophage T4 *td* Group I self-splicing intron and has been engineered and repurposed as a modular, tightly regulated system that can control the expression of any gene of interest (GOI) in a wide range of bacterial species. To illustrate this, we used SIBR to control the Cas12a nuclease from *Francisella novicida* (FnCas12a) and demonstrated efficient genome editing in three bacterial species (*Escherichia coli* MG1655, *Pseudomonas putida* KT2440 and *Flavobacterium* IR1) without the use of exogenous recombinases nor the use of inducible promoters. SIBR is an elegant solution for the widespread problem of engineering prokaryotic organisms with poor recombination efficiencies. We also suggest that SIBR can be used as a universal OFF/ON or ON/OFF switch for individual genes or multiple genes in a polycistronic operon.

## MATERIALS AND METHODS

### Bacterial strains, handling and growth conditions

*E. coli* DH5a (NEB) was used for general plasmid propagation and standard molecular techniques. *E. coli* DH10B T1^R^ (Invitrogen) was used for the LacZ assays. *E. coli* MG1655 (ATCC) was used for targeting and knock-out assays. Unless specified otherwise, *E. coli* strains were grown at 37°C in LB liquid medium (10 g L^-1^ tryptone, 5 g L^-1^ yeast extract, 10 g L^-1^ NaCl) or on LB agar plates (LB liquid medium, 15 g L^-1^ bacteriological agarose) containing the appropriate antibiotics: spectinomycin (100 mg L^-1^), kanamycin (50 mg L^-1^), ampicillin (100 mg L^-1^) or chloramphenicol (35 mg L^-1^). Transformation of electro-competent *E. coli* cells was performed in 2 mm electroporation cuvettes with an ECM 63 electroporator (BTX) at 2500 V, 200 Ω and 25 μF.

*P. putida* strain KT2440 was obtained from DSMZ. Cells were grown at 30°C in LB liquid medium or on LB agar plates containing kanamycin (50 mg L^-1^). Electro-competent *P. putida* cells were transformed in 2 mm electroporation cuvettes using 2500 V, 200 Ω and 25 μF.

*Flavobacterium* species Iridescence 1 (sp. IR1) was kindly provided by Hoekmine BV. *Flavobacterium* IR1 was grown at 25°C in ASW medium (5 g L^-1^ peptone, 1 g L^-1^ yeast extract, 10 g L^-1^ sea salt) or plated on ASW agar (ASW medium, 15 g L^-1^ agar) containing erythromycin (100 mg L^-1^). Electro-competent *Flavobacterium* IR1 cells were transformed in 1 mm electroporation cuvettes using 1500 V, 200 Ω and 25 μF.

### Electro-competent cell preparation

Pre-cultures of *E. coli* or *P. putida* were grown overnight at 37°C in fresh 10 mL LB broth. 5 mL of the overnight culture was inoculated in 500 mL of pre-warmed 2×YP medium (16 g L^-1^ peptone, 10 g L^-1^ yeast extract) and incubated at 37°C shaking at 200 rpm until an OD_600_ of 0.4 was reached. The culture was then cooled down to 4°C. Next, the culture was aliquoted into two sterile 450 mL centrifuge tubes and centrifuged at 3000 g for 10 minutes at 4°C. The supernatant was decanted and the pellet was washed with 250 mL ice-cold sterile miliQ water followed by centrifugation at 3000 g for 10 minutes. The supernatant was decanted and the pellet was resuspended using 5 mL of ice cold 10% glycerol. The two resuspensions were combined in one tube and ice-cold 10% glycerol was added to reach a final volume of 250 mL, followed by centrifugation at 3000 g for 10 minutes. The supernatant was decanted and the pellet was washed with 250 mL of ice-cold 10% glycerol and centrifuged at 3000 g for 10 minutes. The supernatant was decanted and the pellet was resuspended with 2 mL of ice-cold 10% glycerol and aliquoted into tubes of 40 μL. All the electrocompetent cells were stored at - 80°C prior to transformation. To prepare electro-competent cells, *Flavobacterium* IR1 was grown overnight in 10 mL ASW at 25°C, shaking at 200 rpm. The overnight culture was used to inoculate 500 mL ASW in 2 L baffled flask to a starting OD_600_ of 0.05 and incubated at 25°C and shaking at 200 rpm until the cell density reached an OD_600_ equal to 0.3-0.4. The cells were cooled down at 4°C and kept cold on ice for the rest of the procedure. The culture was divided into two sterile 450 mL centrifuge tubes and centrifuged at 3000 g for 10 minutes at 4°C. The supernatant was decanted, and the cell pellet was washed twice with 250 mL ice cold washing buffer (10 mM MgCl_2_ and 5 mM CaCl_2_) followed by centrifugation at 3000 g for 10 min at 4°C. The supernatant was removed, and the pellet was resuspended using 5 mL of the washing buffer. All the resuspensions were combined in one tube and washed by adding 250 mL of the washing buffer. It was then washed once with 10% glycerol followed by centrifugation at 3000 g for 10 minutes. The supernatant was decanted, and the resulting cell pellet was resuspended with 5 mL of ice cold 10% (v/v) glycerol and 100 μL aliquots were stored at −80°C until use.

### Plasmid construction

The LacZ reporter plasmid series were constructed from pEA001 [PWW]. The LacZ reporter plasmid series contain the *E. coli LacZ* gene under the control of the lacUV5 promoter. Ten amino acids flanking the T4 *td* intron (five from each side) were introduced between D6 and S7 of LacZ, omitting the intron itself. For cloning purposes, the ten amino acids were in turn flanked by a BspTI and PstI restriction sites. Generating the complete mutant series was performed by PCR, digestion with BspTI and PstI (Thermo Fisher Scientific) and ligation into pEA001 [PWW]. The main components (origin of replication, antibiotic resistance gene and promoters) of the SIBR-Cas plasmids for *E. coli* and *P. putida* were designed to be functional in both organisms. The LacUV5 promoter was used to drive the expression of the *FnCas12a* variants (WT and Int1-4) and the crRNA. The empty vectors pSIBR001-005 were designed to allow convenient insertion of new spacers through Golden Gate Assembly using the BbsI-HF^®^ enzyme and the T4 DNA ligase (NEB), following the protocol as previously described by Batianis et al. (2019)^13^. Homology arms (500 bp) were introduced to the SIBR-Cas plasmids at a multiple cloning site (MCS). Briefly, homology arms were amplified from genomic DNA and introduced to the MCS of the linearized SIBR-Cas plasmid using the NEBuilder^®^ HiFi DNA Assembly Master Mix (NEB). The plasmid was linearized using Esp3I (NEB). The DNA sequence of all plasmids was verified through Sanger sequencing (Macrogen Europe B.V.). To construct the SIBR-Cas plasmids for *Flavobacterium* IR1, the backbone of pSpyCas9Fb_NT^14^ was used but the Cas9 and the sgRNA were replaced with the *FnCas12a* variants (WT and Int1-4) and the crRNA. Spacers and homology arms were introduced through Golden Gate using BsaI-HF^®^ enzyme and NEBuilder^®^ HiFi DNA Assembly, respectively, as described above. Other plasmids and oligonucleotides used in this study are listed in Supplementary Table 1 and 2 respectively.

### Chemicals and reagents

Unless otherwise specified, all chemical reagents were purchased from Sigma-Aldrich. Sea salt was purchased from Sel Marine. A 40 mM theophylline (Sigma-Aldrich) stock was prepared by dissolving theophylline in dH_2_O followed by filter sterilization using a 0.2 μm Whatman^®^ puradisc syringe filter. When necessary, 0-10 mM theophylline was added to the liquid or solid medium. 20 mg mL^-1^ X-Gal (Sigma-Aldrich) stock was prepared by dissolving X-Gal in N,N-Dimethylmethanamide. The final X-Gal concentration for blue/white colony screening was 0.2 mg mL^-1^.

### β-galactosidase activity assay

LacZ activity was assayed in *E. coli* DH10B T1^R^ in triplicate. After overnight growth at 37°C, 20 μL of culture was mixed with 80 μL of permeabilization solution (100 mM Na_2_HPO_4_, 20 mM KCl, 2 mM MgSO_4_, 0.8 g L^-1^ CTAB, 0.4 g L^-1^ sodium deoxycholate and 5.4 mL L^-1^ β-mercaptoethanol) and incubated at 30°C for 30 min. 600 μL of pre-warmed substrate solution (60 mM Na_2_HPO_4_, 40 mM NaH_2_PO_4_, 1 g L^-1^ o-nitrophenyl-β-D-galactopyranoside and 2.7 mL L^-1^ β-mercaptoethanol) was added and incubated at 30°C until sufficient colour had developed. 700 μL of stop solution (1 M Na_2_CO_3_) was added to quench the reaction. The reaction was filtered through a 0.2 μM filter and measured in a spectrophotometer at 420 nm in a 1 cm cuvette. LacZ activity was calculated according to the following equation:

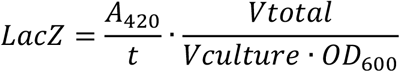

The LacZ activities of all clones were divided by the LacZ activity exhibited by the wild-type intron.

### SIBR-Cas targeting and editing assays in *E. coli*

For both the targeting and the editing assays, 10 ng of plasmid DNA was used to transform 20 μL electro-competent cells as described above. Transformed cells were recovered in 1 mL LB liquid medium for 1 h at 30°C and shaking at 200 rpm. For the targeting assay, the transformants were ten-fold serially diluted in LB liquid medium and 3 μL were used for spot dilution assays on LB agar plates containing kanamycin (50 mg L^-1^) in the presence or absence of 2 mM theophylline and incubated for 24 h at 30°C. For the editing assay, transformants were plated on LB agar plates containing kanamycin (50 mg L^-1^) and 7 mM theophylline and incubated at 30°C for 24 h. Colony PCR was performed on 16 colonies from each transformation to define the editing efficiencies. Triplicate transformations were used for each SIBR-Cas plasmid. Mutant colonies were sequenced through Sanger sequencing (Macrogen BV) to confirm complete deletion of the target gene.

### SIBR-Cas targeting and editing assays *in P. putida*

40 μL electro-competent *P. putida* cells were transformed with 200 ng of plasmid DNA and recovered in 1 mL LB liquid medium for 2 h at 30°C, shaking at 200 rpm. Targeting was assayed by spot dilution assays on LB agar plates containing kanamycin (50 mg L^-1^) in the presence or absence of 2 mM theophylline, followed by overnight incubation at 30°C. *P. putida* cells bearing the editing plasmid were plated on LB agar plates containing kanamycin (50 mg L^-1^) and 2 mM theophylline and incubated at 30°C for 24 h. Grown colonies were screened through colony PCR to define the editing efficiency. For each SIBR-Cas plasmid, transformations were performed in triplicate. Mutant colonies were sequenced through Sanger sequencing (Macrogen BV) to confirm complete deletion of the target gene.

### SIBR-Cas targeting and editing assays *in Flavobacterium* IR1

Electro-competent *Flavobacterium* IR1 cells (100 μL) were transformed with 2 μg plasmid DNA. Transformed cells were recovered in 1 mL ASW and incubated for 4 h at 25°C, shaking at 200 rpm. Due to very low transformation efficiency, the recovered cells were transferred in 10 mL ASW liquid medium containing erythromycin and incubated for 4 d at 25°C, shaking at 200 rpm. 10^-6^ or 10^-7^ cells were then plated on ASW agar containing erythromycin (200 mg L^-1^) and 2 mM theophylline. Plates were incubated at 25°C for 2-3 d and grown colonies were screened for editing through colony PCR. Each editing assay was performed in triplicate. Mutant colonies were sequenced through Sanger sequencing (Macrogen BV) to confirm complete deletion of the target gene.

## RESULTS AND DISCUSSION

### The flanking regions of the T4 *td* intron are amenable to modifications

To create a universal gene control system, we focused on promoter-, sequence- and organism-independent mechanisms. We chose the T4 bacteriophage Group I self-splicing intron that resides in the *thymidylate synthase* (*td*) gene as the appropriate mechanism to control the expression of the GOI (Fig. 1A). The self-splicing ribozyme activity of the T4 *td* intron requires only ubiquitous cofactors such as GTP and Mg^2+^, making its use widely applicable in bacterial species^15^. Moreover, similar to other introns, the T4 *td* intron terminates the translation of the unspliced precursor mRNA due to the presence of in-frame stop codons (Fig. 1B). Therefore, the presence of the intron in the precursor mRNA will result in a truncated non-functional protein, whereas the spliced mRNA allows for the full translation of the protein of interest.

**Figure 1.**
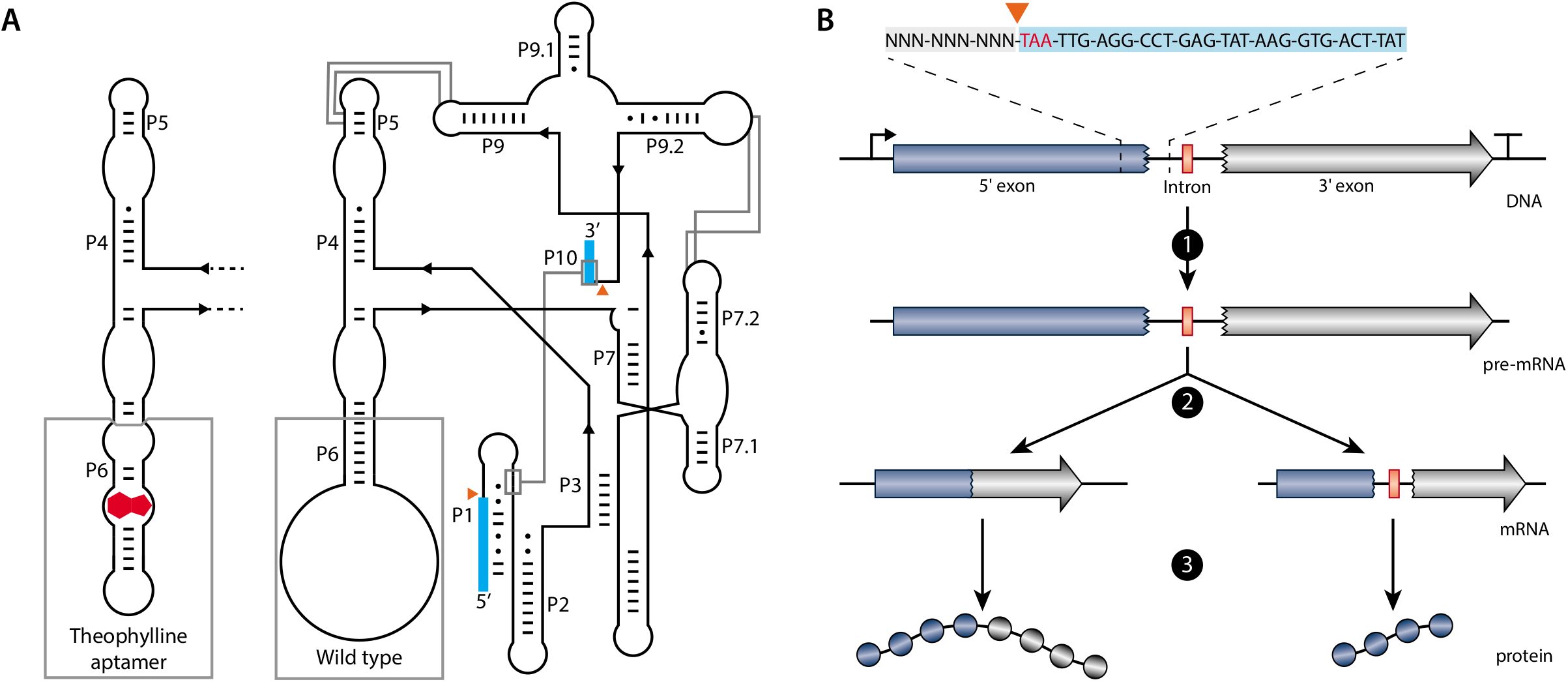
Schematics and function of the T4 *td* intron. (**A**) Schematic representation of the predicted secondary and tertiary structure of the wild type (WT; right) and the theophylline-dependent T4 *td* intron (left). The structure follows the format of Cech et al. (1994)^23^. P1 to P10 represent the pairing domains of the intron. Exon sequences are indicated as blue boxes. Orange triangles indicate the splicing sites. Base pairs are indicated by “-” and wobble pairs by The grey boxes at P6 highlight the difference between the WT and the theophylline-dependent ribozyme. Grey lines show interactions within the intron. (**B**) Schematic representation of the transcription and translation of a gene containing the T4 *td* intron in its open reading frame. At the top, the intron in-frame stop codon (TAA) is depicted in red, the 5’ flanking region is highlighted with a grey box and a part of the intron sequence is highlighted with a light blue box. 1 depicts transcription, 2 depicts self-splicing (left path) or no self-splicing of the intron (right path) and 3 depicts translation of the full protein (left path) or the translation of a truncated protein (right path) when the intron is absent or present, respectively.

The naturally occurring T4 *td* intron is specific for the *td* gene because the exonic flanking regions are necessary to preserve the secondary structure of the P1 and P10 of the T4 *td* intron (Fig. 1A and 2A). Hence, transferring the intron and the flanking exonic regions to another gene will disrupt the coding sequence of the target gene. On the other hand, changing the exonic flanking regions of the intron to preserve the coding sequence of the target gene may affect (or completely inhibit) the splicing activity of the intron. To mitigate this, we opted to create several variants of the T4 *td* intron (with different flanking exons) and placed them in between the coding sequence encoding amino acids D6 and S7 of the *LacZ* gene (Fig. 2A and B). Splicing was assessed through the well-established β-galactosidase assay in *E. coli* DH10B.

**Figure 2.**
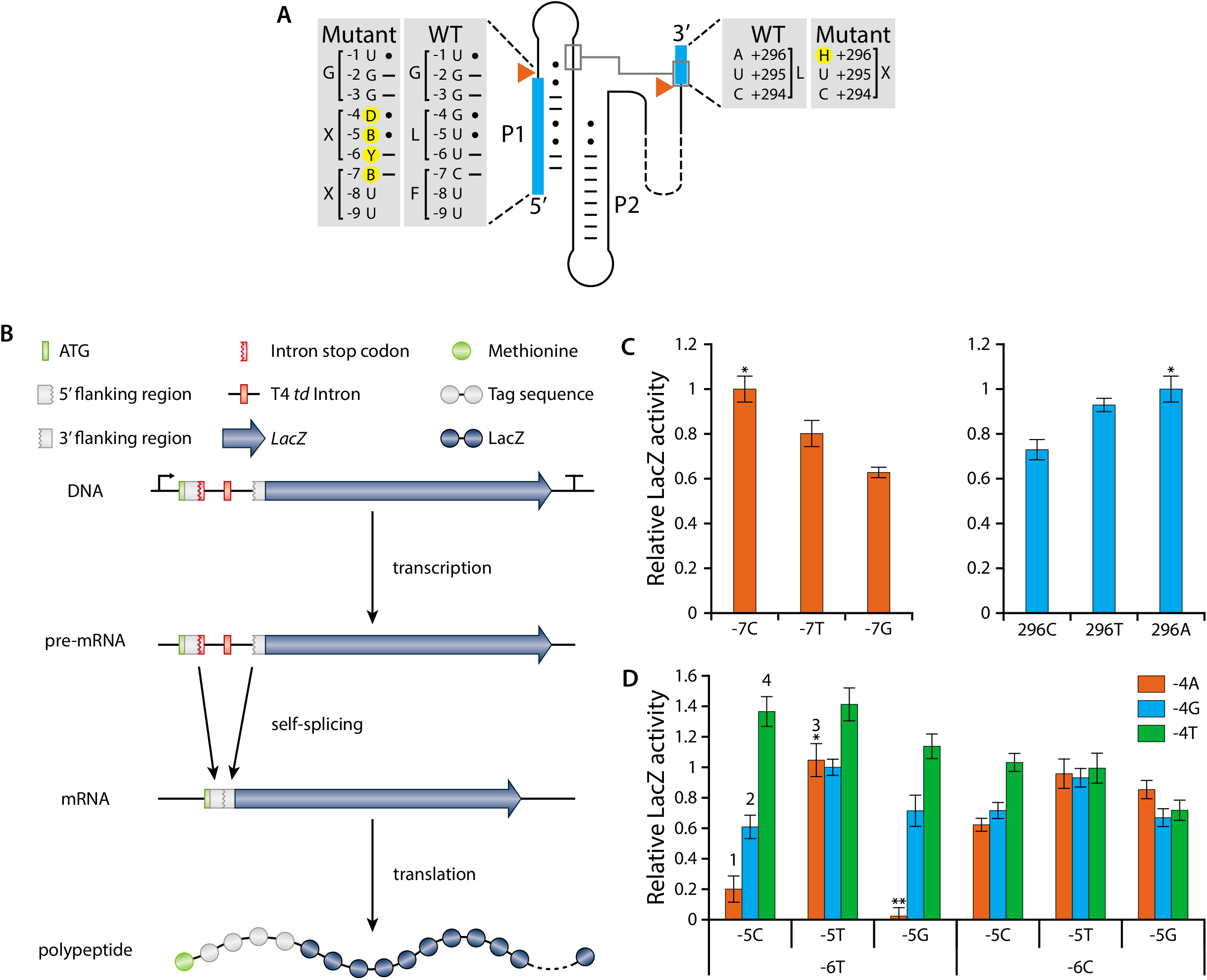
T4 td intron mutant library generation and LacZ assays. (**A**) Detailed illustration of the WT and mutant 5’ and 3’ flanking regions of the T4 *td* intron. Exons are indicated with blue boxes. Orange triangles show the splicing sites. Base pairs are indicated by “-” and wobble pairs by “•”. Mutant nucleotides are highlighted with yellow circles and follow the IUPAC nucleotide nomenclature (B = C/G/T, D = A/G/T, H = A/C/T and Y = C/T). (**B)** *LacZ* transcription-translation cascade controlled by the T4 *td* intron library. (**C**) LacZ activity of position −7 and +296 mutants. “*” indicates the WT intron and is set to 1. All other LacZ activities are a fraction of the wild type activity. (**D**) LacZ activity of all possible combinations for pair, wobble pair and mismatch at positions −6 to −4. “*” indicates the WT intron and is set to 1. All other LacZ activities are a fraction of the wild type activity. “**” indicates the stop codon UGA. The numbers above the bars refer to the intron variants that were selected for the subsequent experiments (Int1, Int2, Int3 and Int4). Bars represent the means and error bars represent the standard deviation of three independent experiments.

Single base modifications at the −7 (C to T or G) or +296 (A to T or C) positions decreased the splicing activity of the intron compared to the wild type (WT) sequence (Fig. 2C). Position −7 (C) preferably pairs with position +15 (G) in the WT intron since a weaker interaction in the form of a wobble base pair (T) or no interaction in the form of a mismatch (G), impeded the splicing of the intron. The opposite was observed for position +296 where a mismatch (A) allows for the highest intron splicing activity. The weak wobble base pair (T) impeded splicing to some extent, while the stronger pair (C) decreased the splicing to a larger extent.

Regardless of the impeded self-splicing of the mutant introns at position −7 and +296, self-splicing was still observed indicating that the flanking regions of the T4 *td* intron are amenable to modifications. To this end, we created pair, wobble or mismatch base substitutions at the −4, −5 and −6 positions and characterized the self-splicing activity of the resulting T4 *td* intron variants (Fig. 2D). For simplicity reasons, the rest of the flanking regions of the T4 *td* intron (including the −7 and +296 positions) were kept the same as the WT sequence.

Surprisingly, several intron variants showed better LacZ activity compared to the WT intron (Fig. 2D). A mismatch at position −4 (T) is preferred in almost all variants, except for those in which both −5 (G) and −6 (C) positions are mismatched too. Compared to the WT intron, a 40% increase in LacZ activity was observed when position −4 was mismatched (T) accompanied by a paired (C) or a wobble paired (T) −5 position and a paired (T) −6 position. A wobble base pair at position −5 (T) negates to a large extent the effect that −4 and −6 have on the splicing. In contrast, a pair (C) or a mismatch (G) at position −5 and depending on −4 and −6 positions, can alter the splicing efficiency from very high to very low. Position −6 in general appears in favour of being paired (T). However, the complete stabilisation of the secondary structure of P1 (−4A, −5C, −6T) is inhibiting splicing almost completely. Mismatching positions −4 to −6 (−4T, −5G, −6C) impedes the splicing very moderately. The −6T, −5G, −4A acts as a negative control as this combination forms a stop codon (UGA), as reflected by the absence of relative LacZ activity using this combination.

Taken together, these results demonstrate that T4 *td* intron variants were generated with a range of splicing efficiencies, allowing for tuneable control over the LacZ protein expression based solely on the intron variant. In addition, the transfer of the T4 *td* intron variants from the *td* gene to the *LacZ* gene demonstrates the flexibility of the intron and its potential use as a universal and tunable gene expression control mechanism.

### SIBR-Cas targeting efficiency is tuneable and inducible

To translate our setup to a CRISPR-Cas engineering context, we tested the ability of the T4 *td* intron variants to control the expression of Cas12a from *Francisella novicida* (FnCas12a). We selected four intron variants with distinct splicing efficiency (Int1: −4A, −5C, −6T; Int2: −4G, −5C, −6T; Int3: −4A, −5T, −6T; Int4: −4T, −5C, −6T; Fig. 2D) and inserted them directly after the start codon of the *FnCas12a* gene (Fig. 3A). According to our design, unspliced precursor mRNAs will result short (5 amino acid) peptides due to the TAA stop codon present at the start of the intron (position +1, +2, +3). In contrast, excision of the intron will result in the full FnCas12a protein fused to a short 4 amino acid tag (SSGL for Int1,2 and 4 or SLGL for Int3) at its N terminus. Furthermore, to make splicing inducible, we added a theophylline aptamer at the P6 stem loop of the T4 *td* intron as previously described^16^, resulting in a new tightly-controlled CRISPR-Cas system, which we named SIBR-Cas (Fig. 3A). Wild-type FnCas12a (WT-FnCas12a, without intron) was used as a reference for comparison to the SIBR-Cas variants. The efficiency of targeting for the SIBR-Cas and WT-FnCas12a variants was assessed by transforming *E. coli* MG1655 cells with plasmids expressing the different Cas12a variants with either a *LacZ* targeting (T) or a non-targeting (NT) crRNA. After transformation, the cells were serially diluted and plated on media with or without the presence of the theophylline inducer (Fig. 3B and).

**Figure 3.**
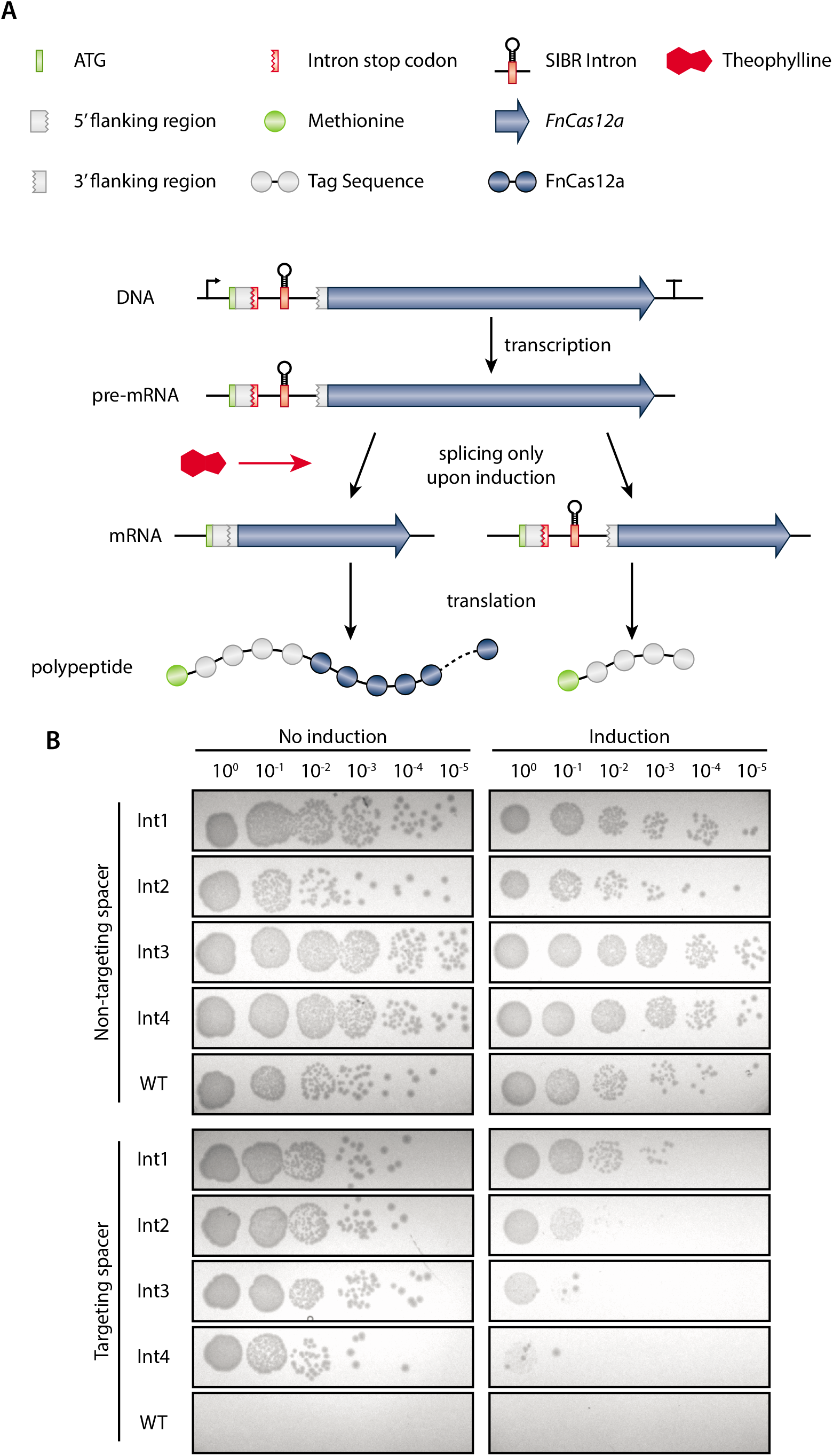
SIBR-Cas targeting assays in *E.coli* MG1655. (**A**) Schematic diagram of SIBR-Cas in the presence or absence of the theophylline inducer. In the presence of theophylline, the intron is self-spliced leading to the translation of the POI with an additional 4 amino acids at its N terminus (left path). In the absence of theophylline, the intron cannot splice itself out of the pre-mRNA, leading to the translation of a short, 5 amino acid long, peptide. (**B**) Targeting and induction efficiency of SIBR-Cas in *E. coli* MG1655. The genome of *E. coli* MG1655 was targeted at the *LacZ* locus with a targeting or a non-targeting spacer. Four intron variants (Int1-4; bad splicer to good splicer) were used to control *FnCas12a* and a WT-FnCas12a was used as a control.

The NT crRNA controls showed colonies up to the 10^-5^ dilution, both in the presence or absence of theophylline for all the SIBR-Cas variants and WT-FnCas12a (Fig. 3B). No colonies were observed when the T crRNA and the WT-FnCas12a combination was used, regardless of induction with theophylline, demonstrating the strong Cas12a-mediated counterselection. In contrast, transformants targeting *LacZ* and expressing either of the four SIBR-Cas variants (Int1-4), showed a notable reduction in colony number formation only when theophylline was present in the medium. Intriguingly, the targeting efficiency directly reflected the splicing efficiency of the intron variants tested for *LacZ* (Fig 2D), with Int1 (−4A, −5C, −6T) showing the least targeting efficiency and Int4 (−4T, −5C, −6T) showing the highest targeting efficiency upon induction. Our results demonstrate consistency in the splicing activity of the intron variants regardless of its genomic context making SIBR a viable strategy for tight, inducible and tuneable control over any GOI.

### SIBR-Cas is an efficient genome engineering tool for bacteria

For efficient genome editing in bacteria, HR should precede CRISPR-Cas counterselection (Fig. 4A). To assess whether tight control over CRISPR-Cas targeting could bolster the efficiency of CRISPR-Cas mediated genome editing by allowing more time for HR to occur, we used SIBR-Cas and targeted the *LacZ* gene of *E. coli* MG1655 for knock-out through HR and CRISPR-Cas counterselection using a blue/white screening colony assay. To facilitate HR, we added 500 bp up- and down-stream homology arms to the plasmids expressing the four SIBR-Cas (Int1-4) and WT-FnCas12a variants that target the *LacZ* gene. After 1 hour recovery, we induced the expression of the SIBR-Cas variants to counterselect the WT from the mutant colonies.

**Figure 4.**
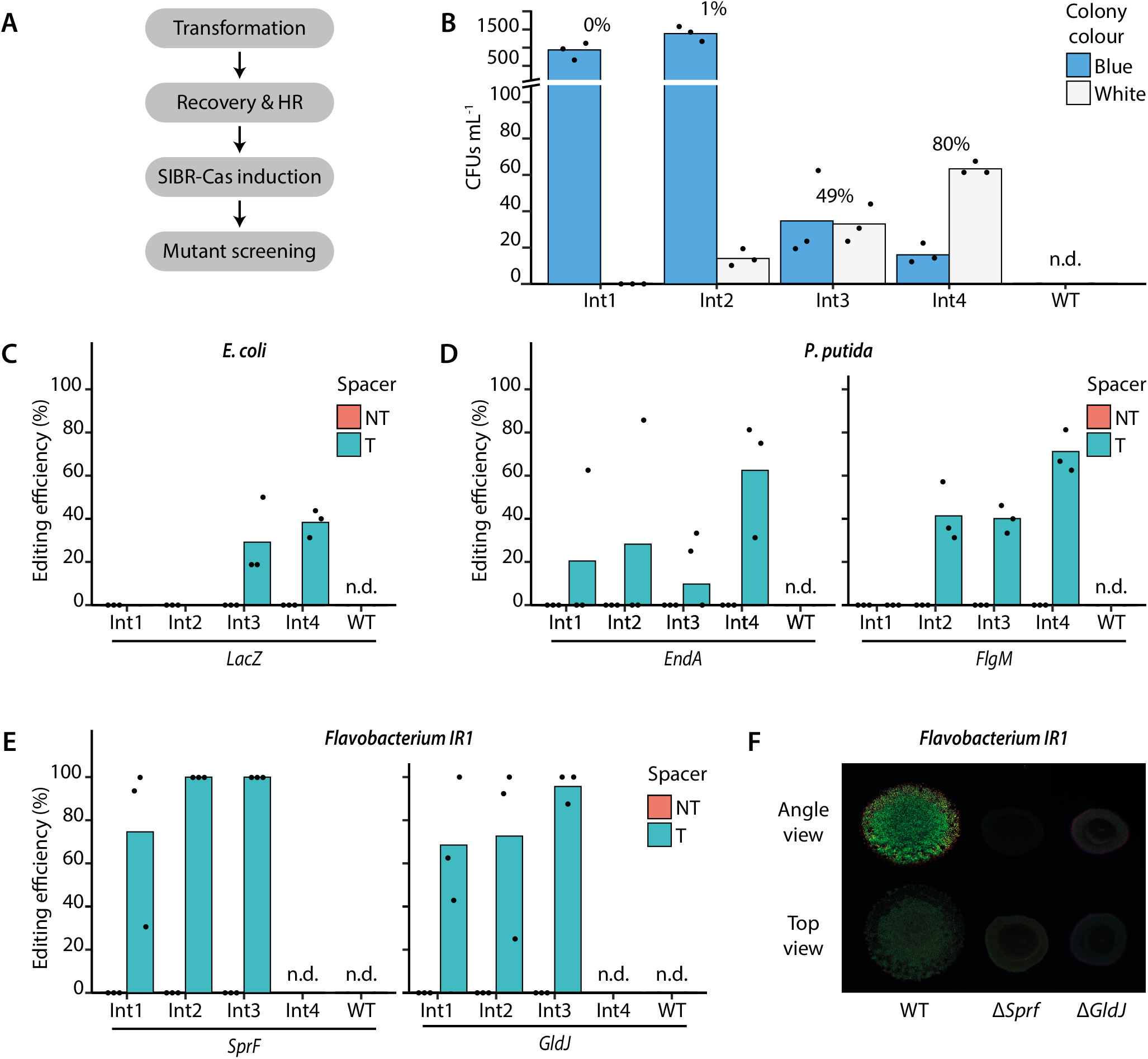
SIBR-Cas genome editing assays in *E. coli* MG1655, *P. putida* KT2440 and *Flavobacterium* IR1. (**A**) Schematic for SIBR-Cas editing procedure. (**B**) Editing efficiency of the *LacZ* gene in *E. coli* MG1655. Blue/white screening was performed to distinguish the edited (white) from the unedited (blue) colonies when using either of the four different SIBR-Cas variants Int1 (0%), Int2 (1% ± 0.35%), Int3 (49% ± 8.75%) and Int4 (80% ± 5.57%) or the WT-FnCas12a (0%). The percentage on top of each variant indicates the percentage of white colonies from the total number of colony forming units mL^-1^ (CFUs mL^-1^). (**C**) Unbiased (omitting the presence of X-Gal in the medium) editing efficiency of the *LacZ* gene using the four different SIBR-Cas variants Int1 (0%), Int2 (0%), Int3 (29% ± 18.04%), Int4 (38% ± 6.41%) or the WT-FnCas12a (0%). (**D**) Editing efficiency of the *EndA* [Int1 (21% ± 36.08), Int2 (29% ± 49.49%), Int3 (19% ± 17.35%), Int4 (63% ± 27.24%) or the WT-FnCas12a (0%)] and *FlgM* [Int1 (0%), Int2 (41% ± 13.84%), Int3 (40% ± 6.41%), Int4 (70% ± 9.85%) or the WT-FnCas12a (0%)] genes in *P. putida* KT2440. (**E**) Editing efficiency of the *SprF* [Int1 (75% ± 38.29), Int2 (100% ± 0%), Int3 (100% ± 0%), Int4 (0%) or the WT-FnCas12a (0%)] and *GldJ* [Int1 (68% ± 29.03%), Int2 (72% ± 41.26%), Int3 (96% ± 7.22%), Int4 (0%) or the WT-FnCas12a (0%)] genes in *Flavobacterium* IR1. Individual bars represent the mean of triplicate experiments and “•” represents the value of each replicate. N.d., not determined. (**F**) Comparison between WT *Flavobacterium* IR1 and Δ*SprF* and Δ*GldJ* strains generated with SIBR-Cas. Images were taken after incubation at room temperature for 2 d by inoculating 3 μL spot on ASWBC medium. The WT strain (left) is 18 mm across.

The WT-FnCas12a variant targeting the *LacZ* gene produced no colonies, demonstrating the targeting efficiency of WT-FnCas12a but also the inefficient HR system of *E. coli* MG1655 (Fig. 4B). In contrast, SIBR-Cas variants produced multiple colonies of which 80% of the total CFUs mL^-1^ were white when Int4 was used, followed by Int3 (49%), Int2 (1%) and Int1 (0%) variants (Fig. 4B and S2). Similar to the previous results, the high editing efficiencies obtained with Int4 suggest that its high splicing efficiency translates into a stronger counterselective pressure. No white colonies were observed for the non-targeting controls, demonstrating that the efficiency of editing without CRISPR-Cas counterselection is negligible (Fig. S3).

Since disruption of LacZ can also be achieved through non-HR mediated approaches (spontaneous mutations or occasional error-prone DNA repair following DNA cleavage by Cas12a), not all gene deletions can be screened phenotypically. Therefore, we repeated our experiment, but X-gal was omitted from the medium to eliminate the possibility of false-positives. Randomly selected colonies that were obtained were screened by PCR for *LacZ* deletion showing a 0%, 0%, 29% and 38% editing efficiency for Int1, Int2, Int3 and Int4 SIBR-Cas variants, respectively (Fig. 4C and S4). The WT-FnCas12a variant targeting *LacZ* did not yield any colonies and all the colonies obtained from the NT controls had the intact, wild-type *LacZ* locus. The observed decrease in editing efficiency (compared to the blue/white screening) might be attributed to spontaneous *LacZ* mutations that escape CRISPR-Cas counterselection. Nevertheless, a high editing efficiency was observed when SIBR-Cas Int4 was used without the use of recombinases or any other complex systems.

Following the successful use of SIBR-Cas in *E. coli*, we continued to demonstrate the universality of SIBR-Cas by testing it in other bacteria. For this purpose, we selected *Pseudomonas putida* KT2440, an organism with rather complex engineering tools and low HR efficiencies^17^. After establishing the successful induction and targeting of SIBR-Cas in *P. putida* (Fig. S5), genome editing experiments were conducted to knock-out the *EndA* and *FlgM* genes. High knock-out efficiencies were obtained when Int4 was used, with editing efficiencies of 63% and 70% for *EndA* and *FlgM*,respectively (Fig. 4D, S6 and S7). Lower editing efficiencies were observed for the other introns, whereas no transformants were obtained with the WT-FnCas12a variant. Control transformants with the NT crRNA had a WT genotype (Fig. S7 and S8).

Lastly, we focused on the non-model organism *Flavobacterium* IR1, which is a recent isolate best known for its iridescent, structural colour^18, 19^. The lack of genomic tools, low transformation efficiency and the low HR efficiency of IR1 are currently the main bottlenecks holding back the fundamental characterization and commercial exploitation of this phenomenon (i.e. development of photonic paints). In addition, *Flavobacterium* species do not have a canonical RBS (TAAAA rather than GGAGG)^20–22^ which render other widely applicable gene control systems, such as Ribo-Cas^8^, inadequate for this type of bacterial species. By using SIBR-Cas, up to 100% efficiencies were achieved for both *SprF* and *GldJ* genes (involved the iridescence) when Int3 was used, whereas Int1 and Int2 showed somewhat lower efficiencies (Fig. 4E, S9 and S10). The WT-FnCas12a variant did not yield any colonies and the nontargeting controls were all confirmed to be unedited (Fig. 4E, S9 and S10). In accordance, the phenotype of the *SprF* mutants displayed similar characteristics when compared to previous studies^18, 19^ (Fig. 4F). Furthermore, SIBR-Cas was successful in creating a clean *GldJ* mutant, that could not be achieved by previous endeavours using transposon mutagenesis (Fig. 4F). Surprisingly and in contrast to *E. coli* and *P. putida*, Int4 failed to sustain growth in the recovery stage (data not shown) and hence was not plated. We expect this to be caused by leakiness of the Int4 variant during the recovery phase for unknown reasons.

Collectively, our results show that SIBR-Cas is a tight and inducible genome engineering tool that can successfully be applied to a wide variety of bacterial species. By delaying CRISPR-Cas counterselection and thus allowing enough time for HR to occur, we achieved high editing efficiencies in model and non-model organisms that naturally have very low HR efficiencies. We propose that this tool could be the solution for the difficulties of using CRISPR-Cas for prokaryotic genome engineering, especially in organisms where HR efficiencies are low, the use of recombinases is not possible or inducible promoters are not characterized. We also foresee that SIBR-Cas will significantly decrease the time required for and complexity of CRISPR-Cas mediated genome engineering in prokaryotes.

### SIBR-X as a modular, tight and inducible protein expression tool

SIBR was successfully applied to control the expression of the *FnCas12a* gene in an OFF to ON manner. We suggest that SIBR-X (where X can be any gene/RNA of interest) can be a broader gene regulation tool for virtually any GOI (Fig. 5A). This is mainly attributed to the host-independent splicing mechanism of the intron variants created during this study. Furthermore, our design of placing the intron directly after the ATG start codon means that it should be compatible with most GOI, leaving only a short four amino acid tag at the N-terminus of the POI, diminishing the risk of interfering with the protein’s functionality. Therefore, with the combination of the intron variants and the theophylline inducer concentration, a temporal and tuneable gene expression can be achieved.

**Figure 5.**
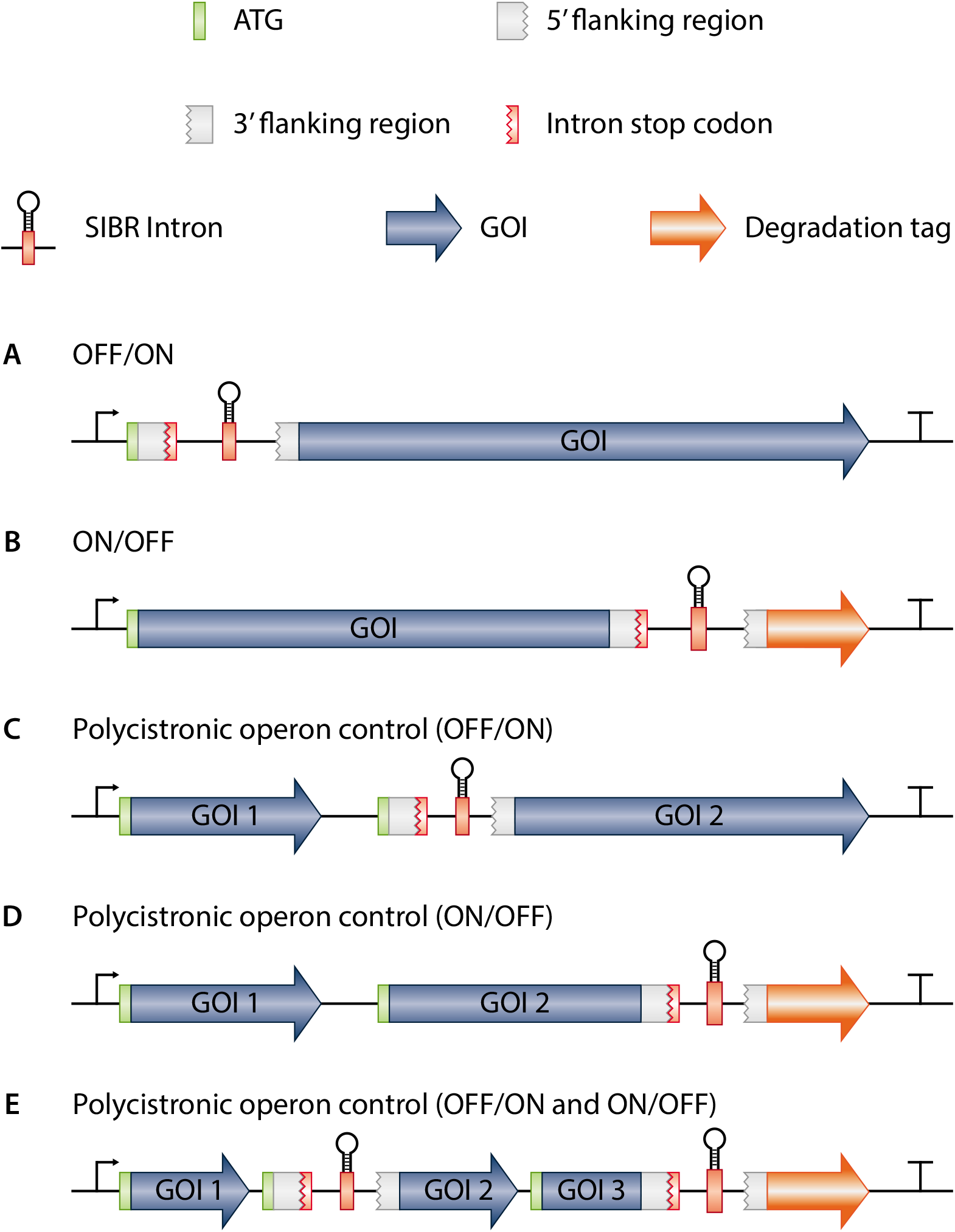
Potential applications of SIBR. (**A**) OFF to ON switch by interrupting the translation of the GOI. (**B**) ON to OFF switch by degrading the POI after inducible attachment of a degradation tag. The degradation tag can be replaced by a localization tag as well. (**C**) Polycistronic operon control allowing constitutive expression of the 1^st^ gene (Gene of interest 1; GOI 1) and inducible OFF to ON expression of the 2^nd^ gene (GOI 2). (**D**) Polycistronic operon control by allowing constitutive expression of both GOI 1 and GOI 2 and inducible protein degradation of GOI 2 upon induction. (**E**) A combination of all the potential SIBR applications in one single polycistronic operon.

SIBR can also be used as an ON to OFF switch (Fig. 5B). For example, SIBR can be inserted at the 3’ of the coding sequence with a downstream degradation tag (e.g. SsrA degradation tag). This design will allow for constitutive translation of the POI in the absence of the inducer (terminated at the stop codon of the intron), but will trigger rapid protein degradation after splicing of the intron due to the attached degradation tag. Other fusions can be envisioned as well, such as a (nuclear) localization tags, signal peptides, etc. Lastly, SIBR can be used as a polycistronic operon control mechanism in different configurations (Fig. 5 C, D and E). This approach will be especially useful in organisms where temporal and inducible expression is difficult to achieve by other means (e.g. operons with multiple and/or uncharacterized promoters and terminators).

Conclusively, we foresee various applications within industry and fundamental research, where SIBR-X can be a valuable tool in both model and non-model organisms.

## ACKNOWLEDGEMENTS

We would like to thank Hoekmine B.V. for sharing their IR1 strain and expertise. We thank Rob Joosten and Steven Aalvink for their technical support.

## Author contribution

C.P., S.C.A.C, S.W.M., J.v.d.O. and R.H.J.S designed the study. C.P., S.C.A.C, A.Q.A and B.A.P. created all the plasmids. S.C.A.C generated and characterized the intron variants. C.P., A.Q.A and B.A.P. generated and characterized SIBR-Cas. C.J.I. provided lab facilities and expertise for handling *Flavobacterium* IR1 through Hoekmine B.V.. C.P., S.C.A.C and R.H.J.S. wrote the manuscript and all authors contributed in reviewing, editing and approving the final manuscript.

