## Supplementary figures and images for "SIBR-Cas enables host-independent and universal CRISPR genome engineering in bacteria"

### Supplemental Figure 1

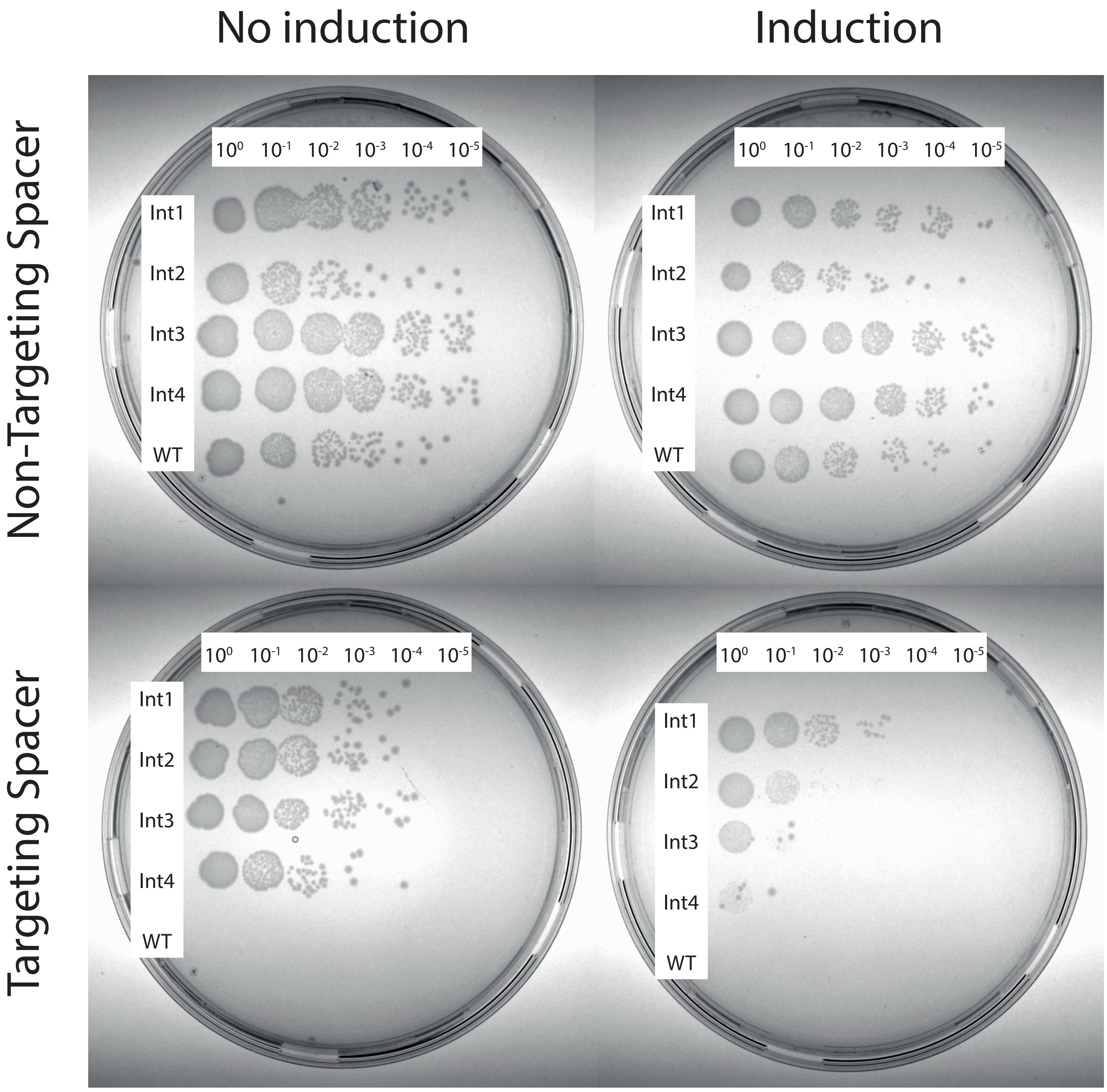

### Supplemental Figure 2

pSIBR016
(Int1)

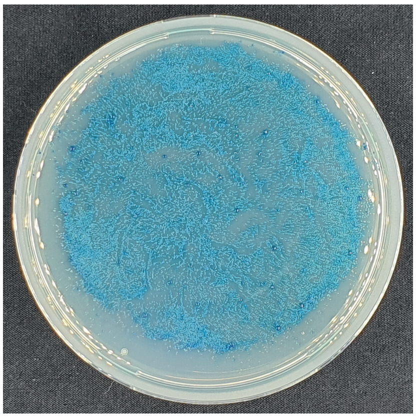

pSIBR017
(Int2)

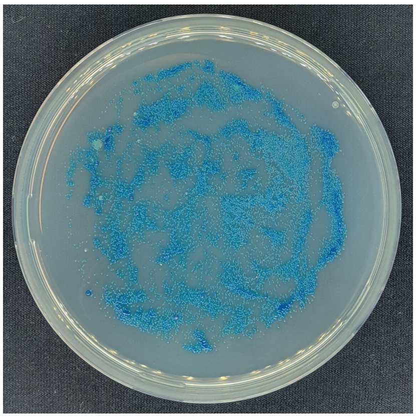

pSIBR018
(Int3)

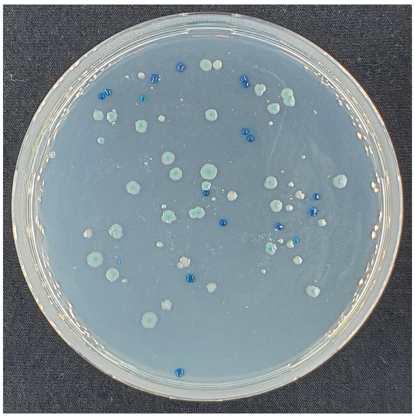

pSIBR019
(Int4)

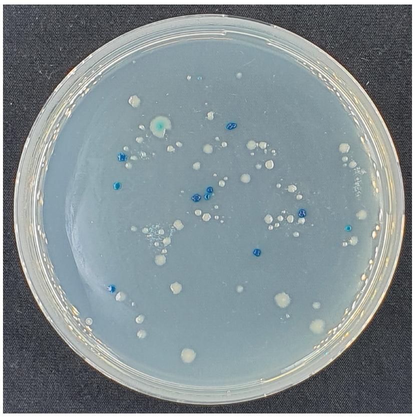

pSIBR009
(WT-FnCas12a)

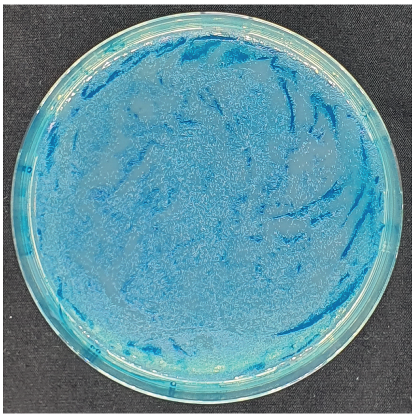

### Supplemental Figure 3

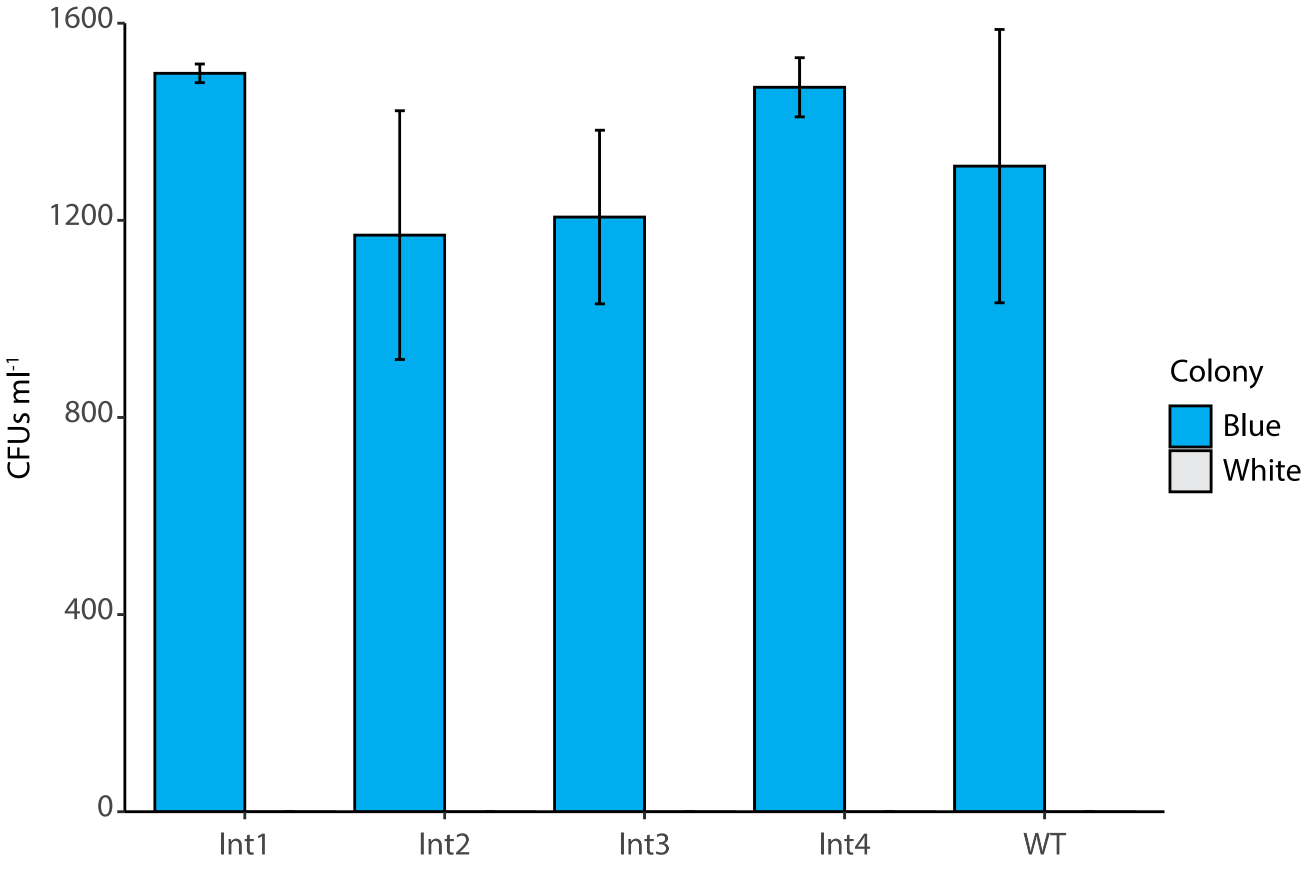

### Supplemental Figure 5

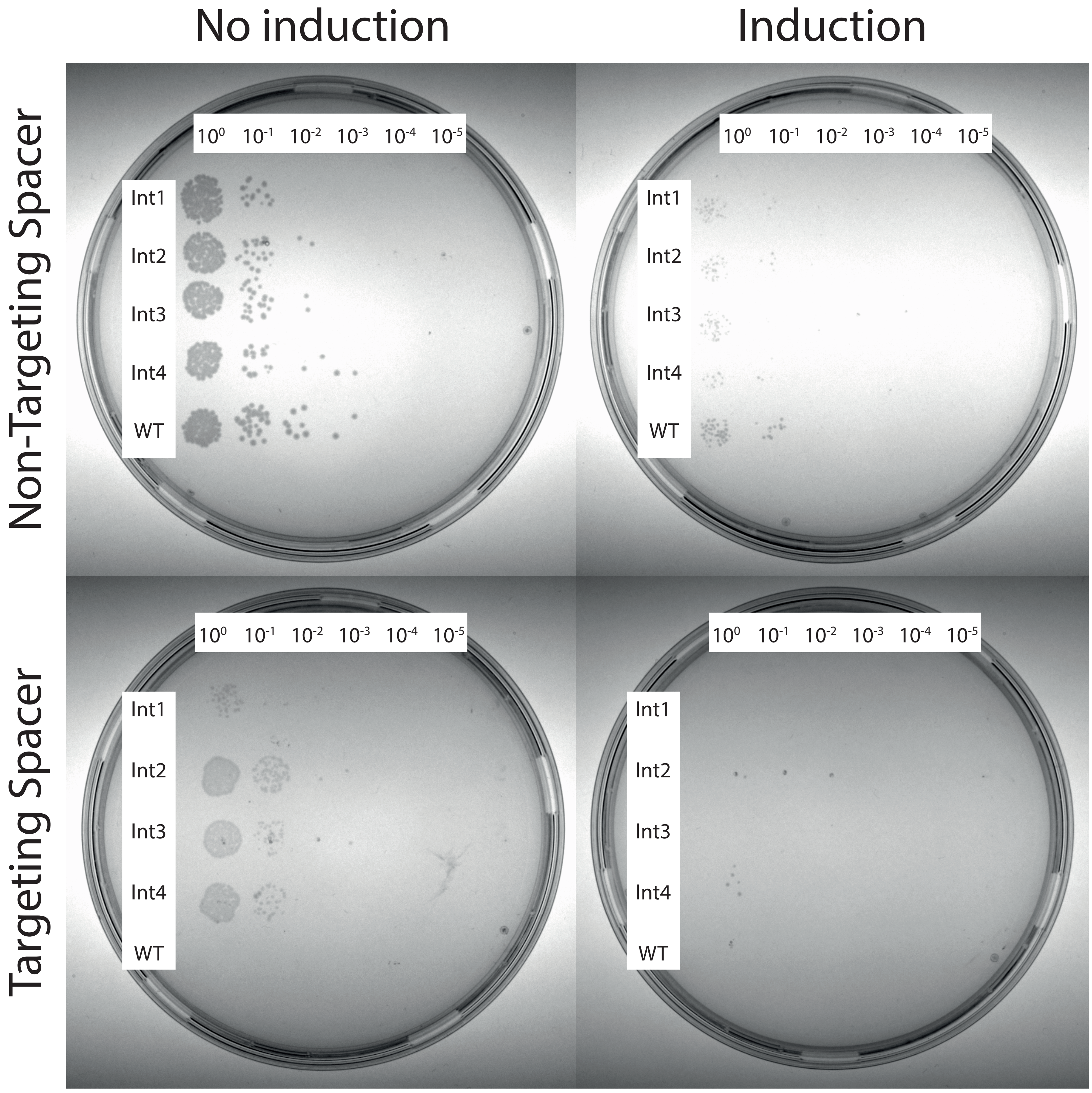

### Supplemental Figure 8

pSIBR024 (Int4-Non Targeting Spacer)

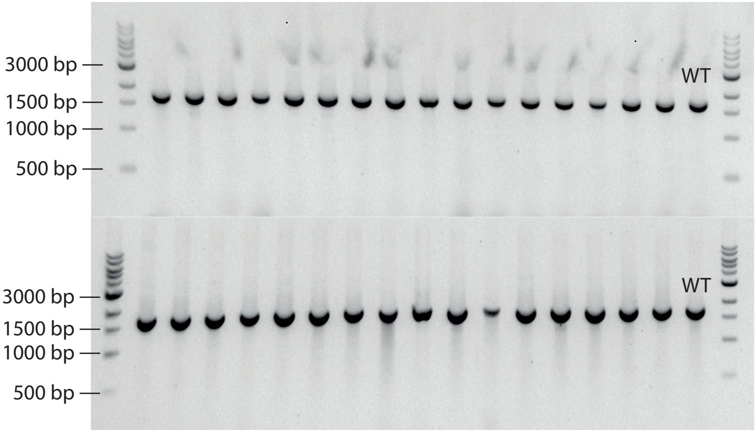
