## Supplemental Figure 4 for "SIBR-Cas enables host-independent and universal CRISPR genome engineering in bacteria"

pSIBR016 (Int1-Targeting Spacer)

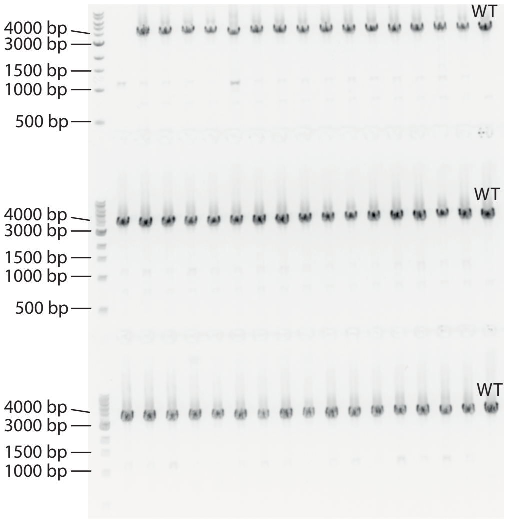


pSIBR017 (Int2-Targeting Spacer)

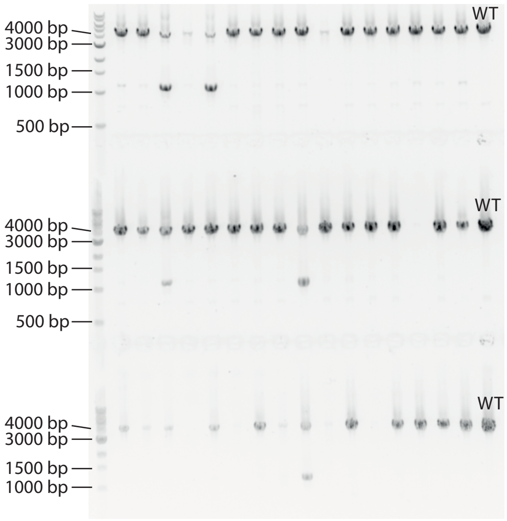


pSIBR018 (Int3-Targeting Spacer)

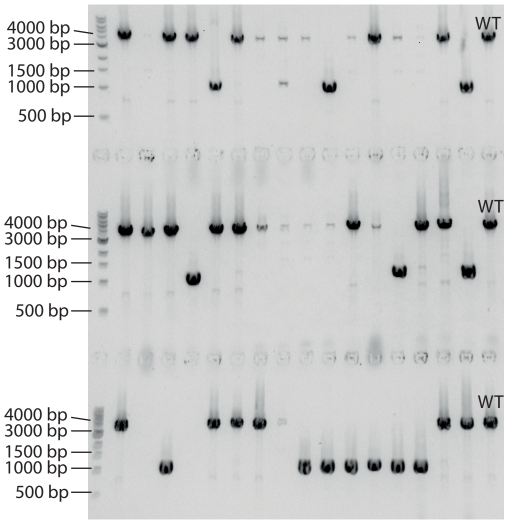


pSIBR019 (Int4-Targeting Spacer)

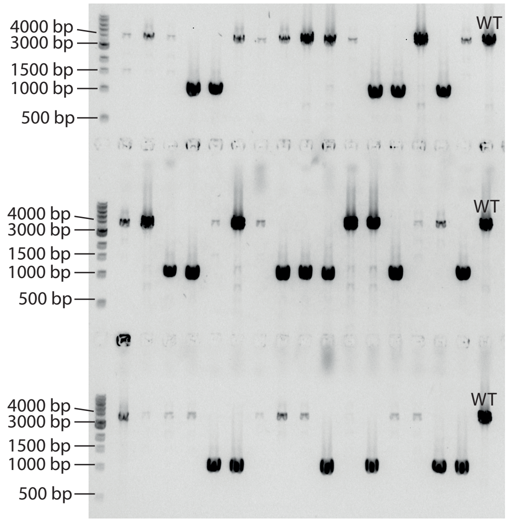


pSIBR009 (Int4-Non Targeting Spacer)

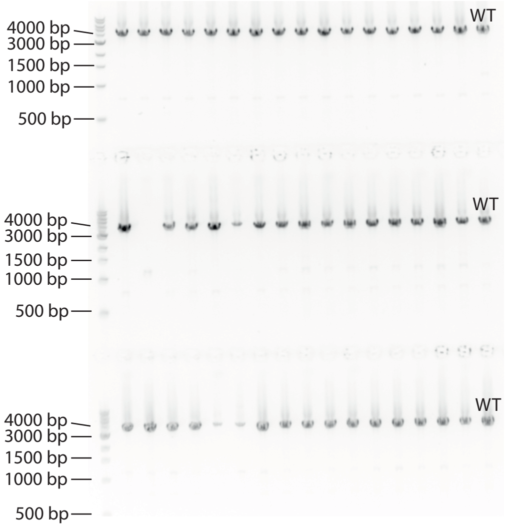
