## Supplemental Figure 6 for "SIBR-Cas enables host-independent and universal CRISPR genome engineering in bacteria"

### pSIBR031 (Int1-Targeting Spacer)

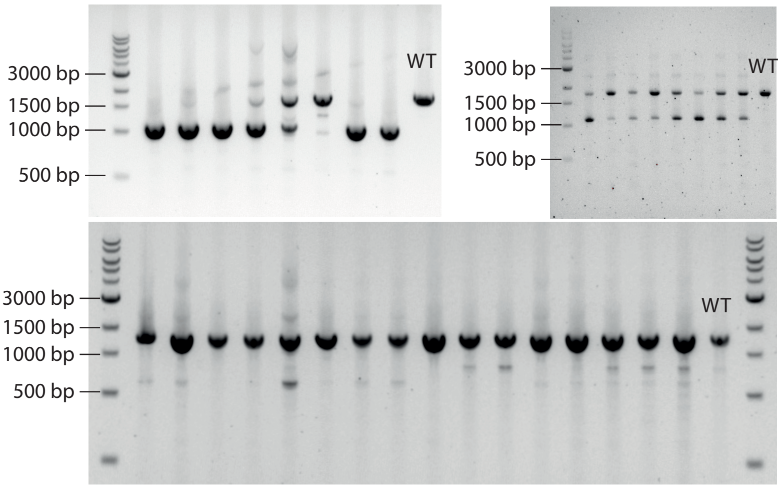


### pSIBR032 (Int2-Targeting Spacer)

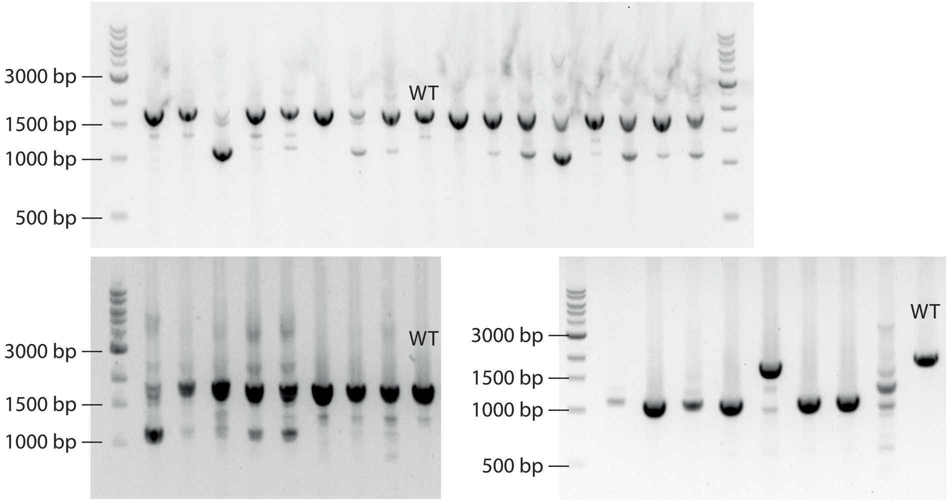


### pSIBR033 (Int3-Targeting Spacer)

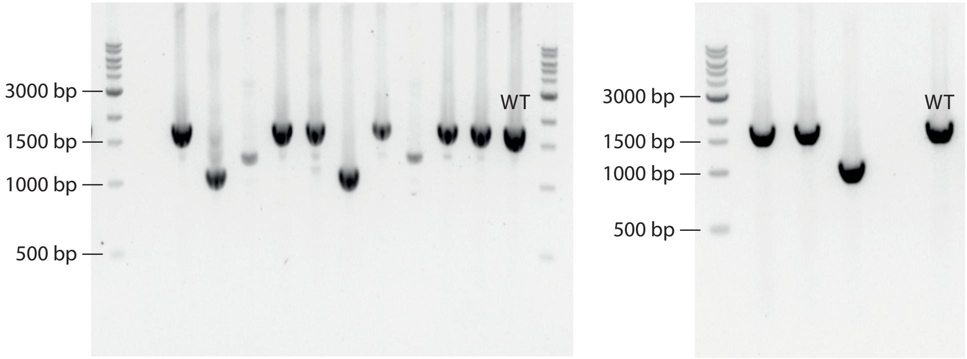


### pSIBR034 (Int4-Targeting Spacer)

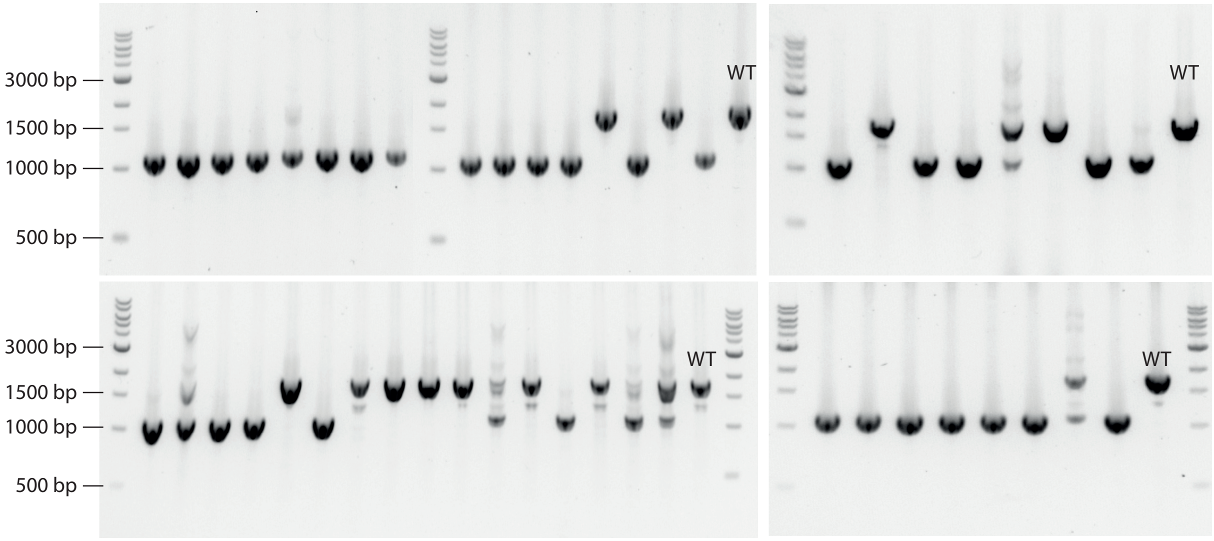
