## Supplemental Figure 7 for "SIBR-Cas enables host-independent and universal CRISPR genome engineering in bacteria"

pSIBR041 (Int1-Targeting Spacer)

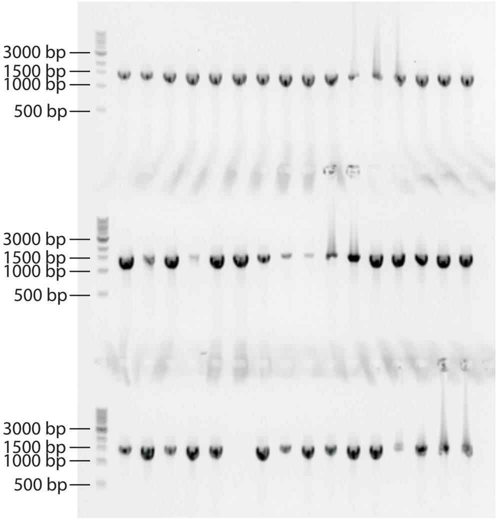


pSIBR042 (Int2-Targeting Spacer)

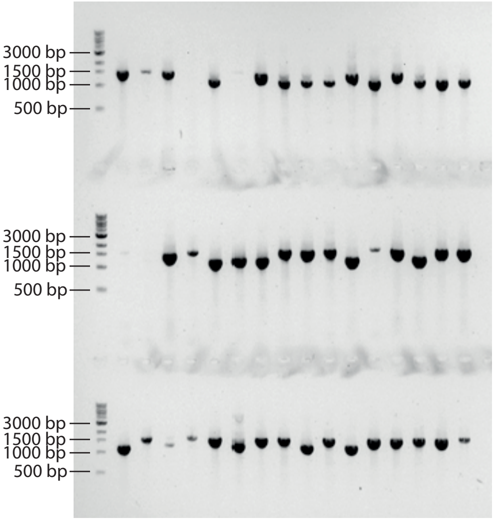


pSIBR043 (Int3-Targeting Spacer)

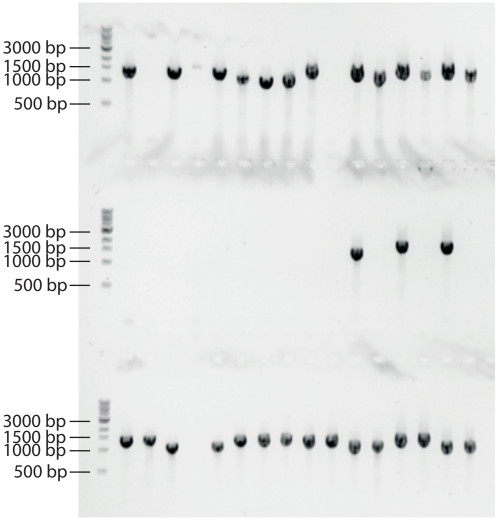


pSIBR044 (Int4-Targeting Spacer)

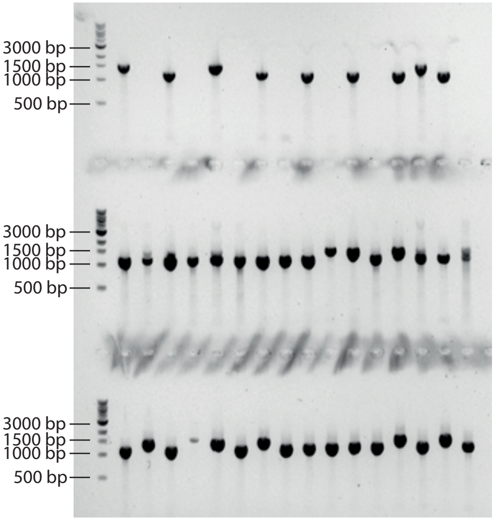


pSIBR039 (Int4-Non Targeting Spacer)

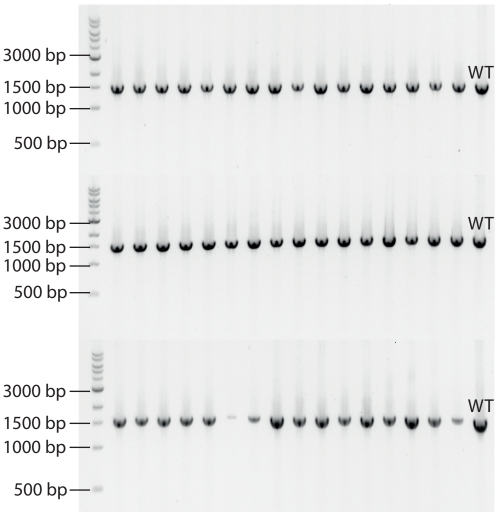
