## Supplemental Figure 9 for "SIBR-Cas enables host-independent and universal CRISPR genome engineering in bacteria"

pSIBR055 (Int1-Targeting Spacer)

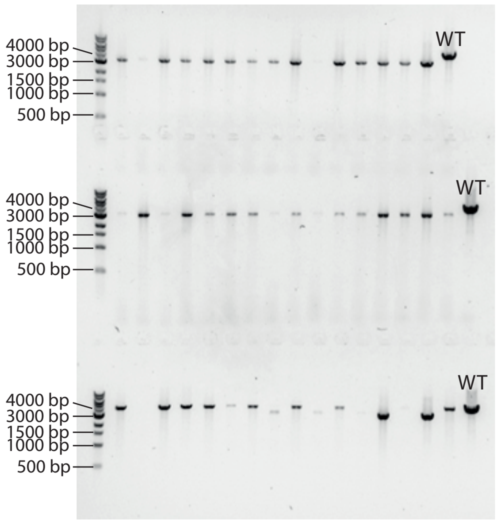


pSIBR056 (Int2-Targeting Spacer)

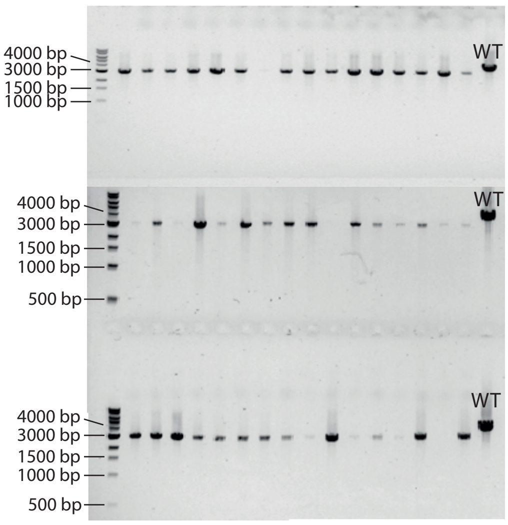


pSIBR057 (Int3-Targeting Spacer)

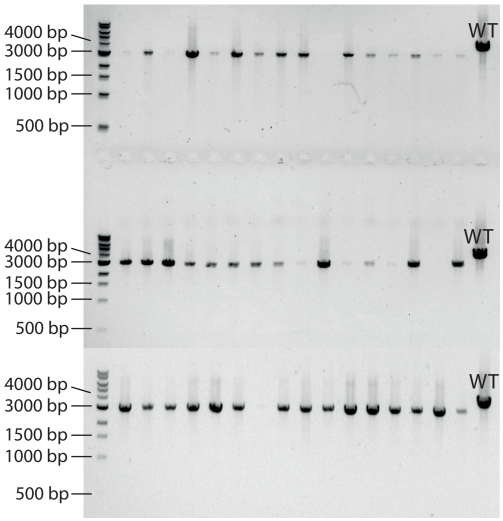


pSIBR053 (Int4-Non Targeting Spacer)

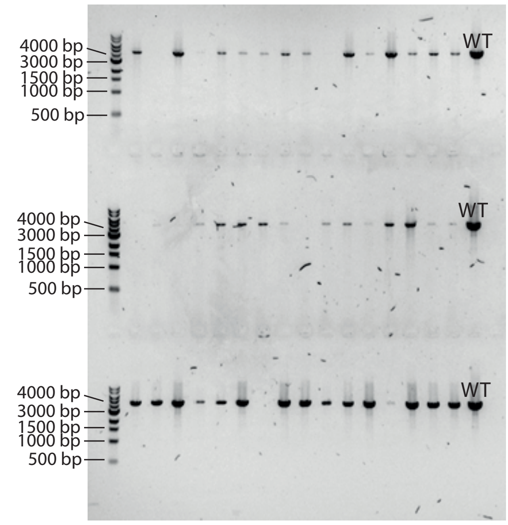
