## Supplemental Figure 10 for "SIBR-Cas enables host-independent and universal CRISPR genome engineering in bacteria"

pSIBR065 (Int1-Targeting Spacer)

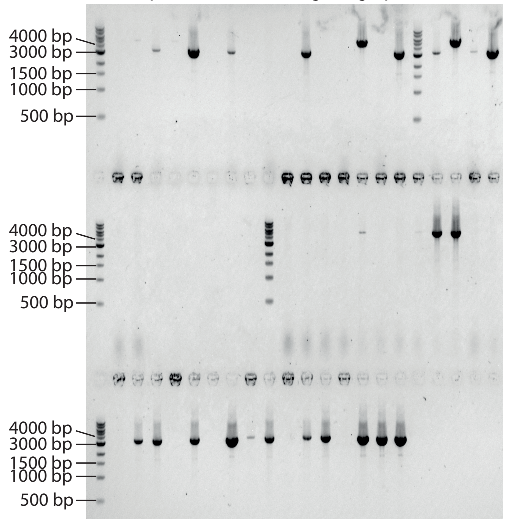


pSIBR066 (Int2-Targeting Spacer)

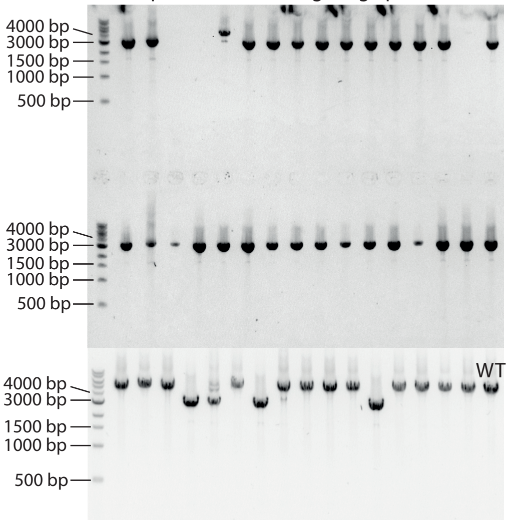


pSIBR067 (Int3-Targeting Spacer)

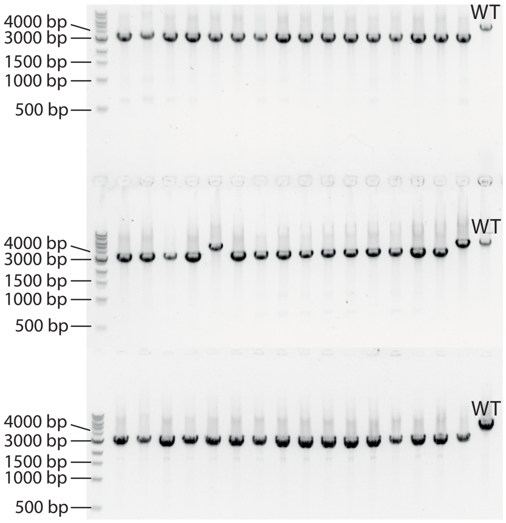


pSIBR063 (Int4-Non Targeting Spacer)

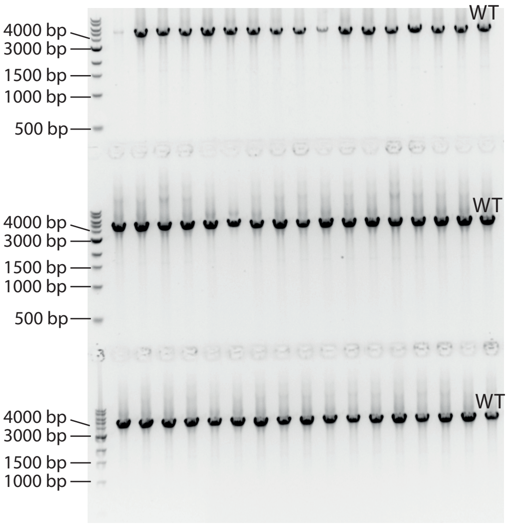
