## Supplemental Figure Legends for "SIBR-Cas enables host-independent and universal CRISPR genome engineering in bacteria"

### SUPPLEMENTARY FIGURE LEGENDS

**Figure S1.** Raw data of SIBR-Cas targeting assay in *E. coli* MG1655. Four different SIBR-Cas variants (Int1-4; Int1 is the worst and Int4 is the best splicer) and the WT-FnCas12a were used for counterselection by targeting *LacZ* with a targeting and a non-targeting spacer. Transformants were serially diluted 5 times and plated on LB medium containing the appropriate antibiotic in the presence or absence of the theophylline inducer (2 mM).

**Figure S2.** Blue/white colony screening of *E. coli* MG1655 transformed with SIBR-Cas plasmids. *E. coli* MG1655 was transformed with plasmids containing either of the four different SIBR-Cas variants (Int1-4; Int1 is the worst and Int4 is the best splicer) or the WT-FnCas12a, *LacZ* targeting or non-targeting spacer and 500 bp homology arms to facilitate the knock-out of *LacZ*. Triplicate experiments were performed. A single representative plate is shown for each triplicate experiment. For the non-targeting variants, only Int4 is shown. WT-FnCas12a transformants targeting *LacZ* did not yield any colonies and are not shown in this figure.

**Figure S3.** Blue/white colony screening of *E. coli* MG1655 transformed with plasmids containing non-targeting spacers. *E. coli* MG1655 was transformed with plasmids containing either of the four different SIBR-Cas variants (Int1-4; Int1 is the worst and Int4 is the best splicer) or the WT-FnCas12a, non-targeting spacer and 500 bp homology arms to facilitate the knock-out of *LacZ*. Values and error bars represent the means and s.d. of triplicate experiments.

**Figure S4.** *LacZ* knock-out using SIBR-Cas. *E. coli* MG1655 was transformed with plasmids containing either of the four different SIBR-Cas variants (Int1-4; Int1 is the worst and Int4 is the best splicer) or the WT-FnCas12a, *LacZ* targeting or non-targeting spacer and 500 bp homology arms to facilitate the knock-out of *LacZ*. Each variant was performed in triplicate and 16 colonies (if present) were randomly selected from each replicate for colony PCR. The WT-FnCas12a variant targeting *LacZ* did not yield any colonies. The non-targeting variants were all WT and for this reason only the Int4 is shown as a representative. Mix amplicons (WT and knock-out bands) were not counted for the total knock-out efficiency percentage. WT: 4229 bp, Knock-out: 1154 bp.

**Figure S5.** Raw data of SIBR-Cas targeting assay in *P. putida* KT2440. Four different SIBR-Cas variants (Int1-4; Int1 is the worst and Int4 is the best splicer) and the WT-FnCas12a were used for counterselection by targeting *EndA* with a targeting and a non-targeting spacer. Transformants were serially diluted 5 times and plated on LB medium containing the appropriate antibiotic in the presence or absence of the theophylline inducer (2 mM).

**Figure S6.** *EndA* knock-out using SIBR-Cas. *P. putida* KT2440 was transformed with plasmids containing either of the four different SIBR-Cas variants (Int1-4; Int1 is the worst and Int4 is the best splicer) or the WT-FnCas12a, *EndA* targeting spacer and 500 bp homology arms to facilitate the knock-out of *EndA*. Each variant was performed in triplicate and 16 colonies (if present) were randomly selected from each replicate for colony PCR. The WT-FnCas12a variant targeting *EndA* did not yield any colonies. Mix amplicons (WT and knock-out bands) were not counted for the total knock-out efficiency percentage. WT: 1814 bp, Knock-out: 1121 bp.

**Figure S7.** *FlgM* knock-out using SIBR-Cas. *P. putida* KT2440 was transformed with plasmids containing either of the four different SIBR-Cas variants (Int1-4; Int1 is the worst and Int4 is the best splicer) or the WT-FnCas12a, *FlgM* targeting or non-targeting spacer and 500 bp homology arms to facilitate the knock-out of *FlgM*. Each variant was performed in triplicate and 16 colonies (if present) were randomly selected from each replicate for colony PCR. The WT-FnCas12a variant targeting *LacZ* did not yield any colonies. The non-targeting variants were all WT and for this reason only the Int4 is shown as a representative. Mix amplicons (WT and knock-out bands) were not counted for the total knock-out efficiency percentage. WT: 1523 bp, Knock-out: 1208 bp.

**Figure S8.** *EndA* knock-out using non-targeting SIBR-Cas. *P. putida* KT2440 was transformed with plasmids containing either of the four different SIBR-Cas variants (Int1-4; Int1 is the worst and Int4 is the best splicer) or the WT-FnCas12a, *EndA* non-targeting spacer and 500 bp homology arms to facilitate the knock-out of *EndA*. Each variant was performed in triplicate and 16 colonies (if present) were randomly selected from each replicate for colony PCR. All variants were WT and for this reason only the Int4 is shown as a representative. WT: 1814 bp, Knock-out: 1121 bp.

**Figure S9.** *SprF* knock-out using SIBR-Cas. *Flavobaterium* IR1 was transformed with plasmids containing either of the four different SIBR-Cas variants (Int1-4; Int1 is the worst and Int4 is the best splicer) or the WT-FnCas12a, *SprF* targeting or non-targeting spacer and 500 bp homology arms to facilitate the knock-out of *SprF*. Each variant was performed in triplicate and 16 colonies (if present) were randomly selected from each replicate for colony PCR. The Int4 and WT-FnCas12a variants targeting *LacZ* did not yield any colonies. The non-targeting variants were all WT and for this reason only the Int4 is shown as a representative. Mix amplicons (WT and knock-out bands) were not counted for the total knock-out efficiency percentage. WT: 4081 bp, Knock-out: 3082 bp.

**Figure S10.** *GldJ* knock-out using SIBR-Cas. *Flavobaterium* IR1 was transformed with plasmids containing either of the four different SIBR-Cas variants (Int1-4; Int1 is the worst and Int4 is the best splicer) or the WT-FnCas12a, *GldJ* targeting or non-targeting spacer and 500 bp homology arms to facilitate the knock-out of *GldJ*. Each variant was performed in triplicate and 16 colonies (if present) were randomly selected from each replicate for colony PCR. The Int4 and WT-FnCas12a variants targeting *LacZ* did not yield any colonies. The non-targeting variants were all WT and for this reason only the Int4 is shown as a representative. Mix amplicons (WT and knock-out bands) were not counted for the total knock-out efficiency percentage. WT: 4944 bp, Knock-out: 3255 bp.
